# CRISPR system acquisition and evolution of an obligate intracellular *Chlamydia*-related bacterium

**DOI:** 10.1101/028977

**Authors:** Claire Bertelli, Ousmane Cissé, Brigida Rusconi, Carole Kebbi-Beghdadi, Antony Croxatto, Alexander Goesmann, François Collyn, Gilbert Greub

**Author notes:** Corresponding author: Gilbert Greub, MD PhD Institute of Microbiology University of Lausanne 1011 Lausanne SWITZERLAND.

## Abstract

Recently, a new *Chlamydia-related* organism, *Protochlamydia naegleriophila* KNic, was discovered within a *Naegleria* amoeba. To decipher the mechanisms at play in the modeling of genomes from the *Protochlamydia* genus, we sequenced *de novo* the full genome of *Pr. naegleriophila* combining the advantages of two second-generation sequencing technologies. The assembled complete genome comprises a 2,885,111 bp chromosome and a 145,285 bp megaplasmid. For the first time within the *Chlamydiales* order, a CRISPR system, the immune system of bacteria, was discovered on the chromosome. It is composed of a small CRISPR locus comprising eight repeats and the associated *cas* and *cse* genes of the subtype I-E. A CRISPR locus was also found within *Chlamydia* sp. Diamant, another *Pr. naegleriophila* strain whose genome was recently released, suggesting that the CRISPR system was acquired by a common ancestor of these two members of *Pr. naegleriophila,* after the divergence from *Pr. amoebophila.* The plasmid encodes an F-type conjugative system similar to that found in the Pam100G genomic island of *Pr. amoebophila* suggesting an acquisition of this conjugative system before the divergence of both *Protochlamydia* species and the integration of a putative *Pr. amoebophila* plasmid into its main chromosome giving rise to the Pam100G genomic island. Overall, this new *Pr. naegleriophila* genome sequence enables to investigate further the dynamic processes shaping the genomes of *Chlamydia-related* bacteria.

## INTRODUCTION

A large diversity prevails in the order *Chlamydiales,* as suggested by the discovery of a large number of *Chlamydia* and *Chlamydia-related* bacteria belonging to nine different families (Greub 2010; Everett et al. 1999; Horn 2011) and the cross-examination of metagenomics data (Lagkouvardos et al. 2014). The family *Parachlamydiaceae* comprises five genera that are each represented by a small number of isolated strains. The genus *Protochlamydia* was lately enriched by the isolation of a *Naegleria* endosymbiont that presented 97.6% identity in the 16S rRNA with *Pr. amoebophila* UWE25 and was thus named *Pr. naegleriophila* strain KNic (Casson et al. 2008). Since other members of the *Parachlamydiaceae* family were suspected to be associated with lung infections (Greub 2009), a diagnostic PCR specific for *Pr. naegleriophila* was developed and applied to bronchoalveolar lavages. *Pr. naegleriophila* DNA was detected in the bronchoalveolar lavage of an immunocompromised patient with pneumonia by two PCRs targeting different genomic regions and the presence of the bacterium in the sample was confirmed by direct immunofluorescence (Casson et al. 2008). These results indicated a potential role of *Pr. naegleriophila* in lower respiratory tract infections.

A recent study including *Chlamydia* genomes and other members of the *Planctomycetes-Verrucomicrobia-Chlamydia* superphylum suggested that the branch leading to *Chlamydia* was shaped mainly by genome reduction and evidenced limited occurrence of gene birth, duplication and transfer (Kamneva et al. 2012), as it is the case in other strict intracellular pathogens (Darby et al. 2007). On the contrary, the occurrence of large families of paralogs in the genome of *Chlamydia-*related bacteria suggested an evolution by extensive gene duplication (Domman et al. 2014; Eugster et al. 2007). The chromosome sequence of *Pr. amoebophila* UWE25 exhibited little evidence for the occurrence of lateral gene transfer (Horn et al. 2004). However, a number of probable lateral gene transfers were identified between *Parachlamydia* and other amoeba-infecting bacteria such as *Legionella* (Gimenez et al. 2011), a process that may take place within the amoeba itself (Bertelli & Greub 2012). The *Pr. amoebophila* genome presents a genomic island (Pam100G) that encodes a type IV secretion system of the F-type that might be involved in conjugative DNA transfer (Greub et al. 2004). A similar system was found on the plasmid of *Simkania negevensis* (Collingro et al. 2011) and a partial operon was described in *Parachlamydia acanthamoebae* (Greub et al. 2009), suggesting active DNA transfer capabilities in the ancestor of the *Chlamydiales* and some of its descendants.

Small interspaced repetitions were initially observed in *E. coli* (Ishino et al. 1987) and they were then named CRISPR, an acronym for Clustered Regularly Interspaced Short Palindromic Repeats (Jansen et al. 2002). Although found in 50% of bacteria and in 90% of archaea (Weinberger et al. 2012), a CRISPR system had never been previously reported in a member of the order *Chlamydiales* (Makarova et al. 2011). The CRISPR locus usually consists of a variable number (up to 587) of 23-47bp repeats with some dyad symmetry, but not truly palindromic, interspaced by 21-72 bp spacers (Horvath & Barrangou 2010). Associated with these repeats are 2 core cas genes and additional subtype-specific genes putatively providing mechanistic specificity (Koonin & Makarova 2013). Similarity between spacers and extrachromosomal elements first suggested a role in immunity against phage infection and more generally against conjugation or transformation by acquisition of external DNA (Bolotin et al. 2005). The CRISPR-Cas system was shown to mediate an antiviral response thus inducing resistance to phage infection (Deveau et al. 2010), notably in *E. coli* (Brouns et al. 2008). More recently, CRISPR-Cas systems were shown to regulate stress-related response, changing gene expression and virulence traits in several pathogens among which is the intracellular bacteria *Francisella novicida* (Sampson & Weiss 2014; Louwen et al. 2014).

In this contribution, we sequenced and analyzed the complete genome of *Pr. naegleriophila* strain KNic and discovered two potentially antagonistic systems, a type IV secretion system likely implicated in conjugative DNA transfer and a CRISPR system that generally controls foreign DNA acquisition. Furthermore, the complete genome sequence of a new species within the genus *Protochlamydia* offered the possibility to look into the genome dynamics throughout evolution by comparing *Pr. naegleriophila* KNic gene content and genome architecture to its closest relatives of the family *Parachlamydiaceae*.

## RESULTS

### Chromosome features and evolution

*Pr. naegleriophila* KNic possesses a 2’885’090 bp circular chromosome with a mean GC content of 42.7%. The genome size and the GC content are surprisingly high compared to the most closely-related species, *Pr. amoebophila* (**Table 1**), but it is consistent with its closest relative *Chlamydia sp.* Diamant, another *Pr. naegleriophila* strain (hereafter referred to as *Pr. naegleriophila* Diamant). The chromosome of *Pr. naegleriophila* strain KNic is predicted to encode 2,415 proteins and exhibits four ribosomal operons and 43 tRNAs, more than any other *Chlamydiales* (**Table 1**). Two types of spacers are found between the 16S and the 23S rRNA: either a simple intergenic spacer or a spacer containing two tRNAs for Ala and Ile.

**Table 1.** Genomics characteristics of bacteria belonging to the family *Parachlamydiaceae*. As available on NCBI database on 22.09.2015, all genomes except KNic have been reannotated by the NCBI Prokaryotic Genome Annotation Pipeline.

| Species | Strain | Status | Scaffolds | Genome size | GC content | % coding | CDS | tRNAs | rRNA genes | Plasmid size | Plasmid CDS | Plasmid GC content |
| --- | --- | --- | --- | --- | --- | --- | --- | --- | --- | --- | --- | --- |
| *Protochlamydia amoebophila* | UWE25 | Complete | 1 | 2,414,465 | 34.7 |  | 1855 | 35 | 7 | - |  |  |
| *Protochlamydia amoebophila* | EI2 | Draft | 178 | 2,397,675 | 34.8 |  | 1797 | 36 | 3 | NA |  |  |
| *Protochlamydia amoebophila* | R18 | Draft | 795 | 2,881,499 | 34.8 |  | 2025 | 41 | 13 | NA |  |  |
| *Protochlamydia naegleriophila* | KNic | Complete | 1 | 2,885,090 | 42.7 |  | 2415 | 43 | 12 | 145,285 | 160 | 37.2 |
| *Protochlamydia naegleriophila* | Diamant | Draft | 4* | 2,864,073 | 42.8 |  | 2424 | 39 | 7 | 91,928 | 98 | 40.9 |
| *Parachlamydia acanthamoebae* | UV7 | Complete | 1 | 3,072,383 | 39 |  | 2531 | 40 | 10 | - |  |  |
| *Parachlamydia acanthamoebae* | Hall's coccus | Draft | 95 | 2,971,261 | 39 |  | 2474 | 35 | 3 | NA |  |  |
| *Parachlamydia acanthamoebae* | OEW1 | Draft | 162 | 3,008,885 | 39 |  | 2321 | 38 | 4 | NA |  |  |
| *Parachlamydia acanthamoebae* | Bn9 | Draft | 72 | 2,999,361 | 38.9 |  | 2498° | NA | NA | NA |  |  |
| *Parachlamydiaceae* bacterium | HS-T3 | Draft | 34 | 2,307,885 | 38.7 |  | 2003 | 39 | 3 | NA |  |  |
| *Parachlamydia* sp. | Rubis | Draft | 3* | 2,701,449 | 32.4 |  | 2446 | 36 | 5 | 80,697° | 107 | 40.2 |
|  |  |  |  |  |  |  |  |  |  | 39,075° | 40 | 29.8 |
| *Neochlamydia* sp. | EPS4 | Draft | 112 | 2,530,677 | 38.1 |  | 1843 | 36 | 4 | NA |  |  |
| *Neochlamydia* sp. | TUME1 | Draft | 254 | 2,546,323 | 38 |  | 1834 | 36 | 4 | NA |  |  |
| *Neochlamydia* sp. | S13 | Draft | 1342 | 3,187,074 | 38 |  | 2175 | 42 | 10 | NA |  |  |

\* Following removal of the plasmid(s) present according to our analyses. ° Not circularized, estimated size . NA : information not available

The cumulative G versus C nucleotide bias (GC skew) presents a typical pyramidal shape (**Figure S1**) that is expected in the absence of particular large genomic islands and confirms the assembly accuracy. The GC skew of *Pr. naegleriophila* is smoother than that of *Pr. amoebophila*, and does not present the small inversion in the slope that is caused by the *Pr. amoebophila* genomic island (**Figure S1**) (Greub et al. 2004). The replication origin *(ori)* and the terminus of replication (*ter*), at the minimum and maximum of the curve (**Figure S1**), respectively, show an almost perfectly balanced chromosome with 49.8% of the base on one arm, i.e., between *ori* and *ter*, and 50.2% on the other arm, i.e., between *ter* and *ori*.

The two strains of *Pr. naegleriophila* are highly collinear as shown in the alignment of available complete and nearly complete (<5 contigs) genomes of the family *Parachlamydiaceae* (**Figure 1**). Within genus comparison of *Pr. naegleriophila* and *Pr. amoebophila* shows the occurrence of 13 recombination and inversion events. As expected, further distantly-related organisms from a different genus exhibit a lower collinearity and an increasing number of recombination events, which is correlated to the phylogenetic distance.

**Figure 1.**
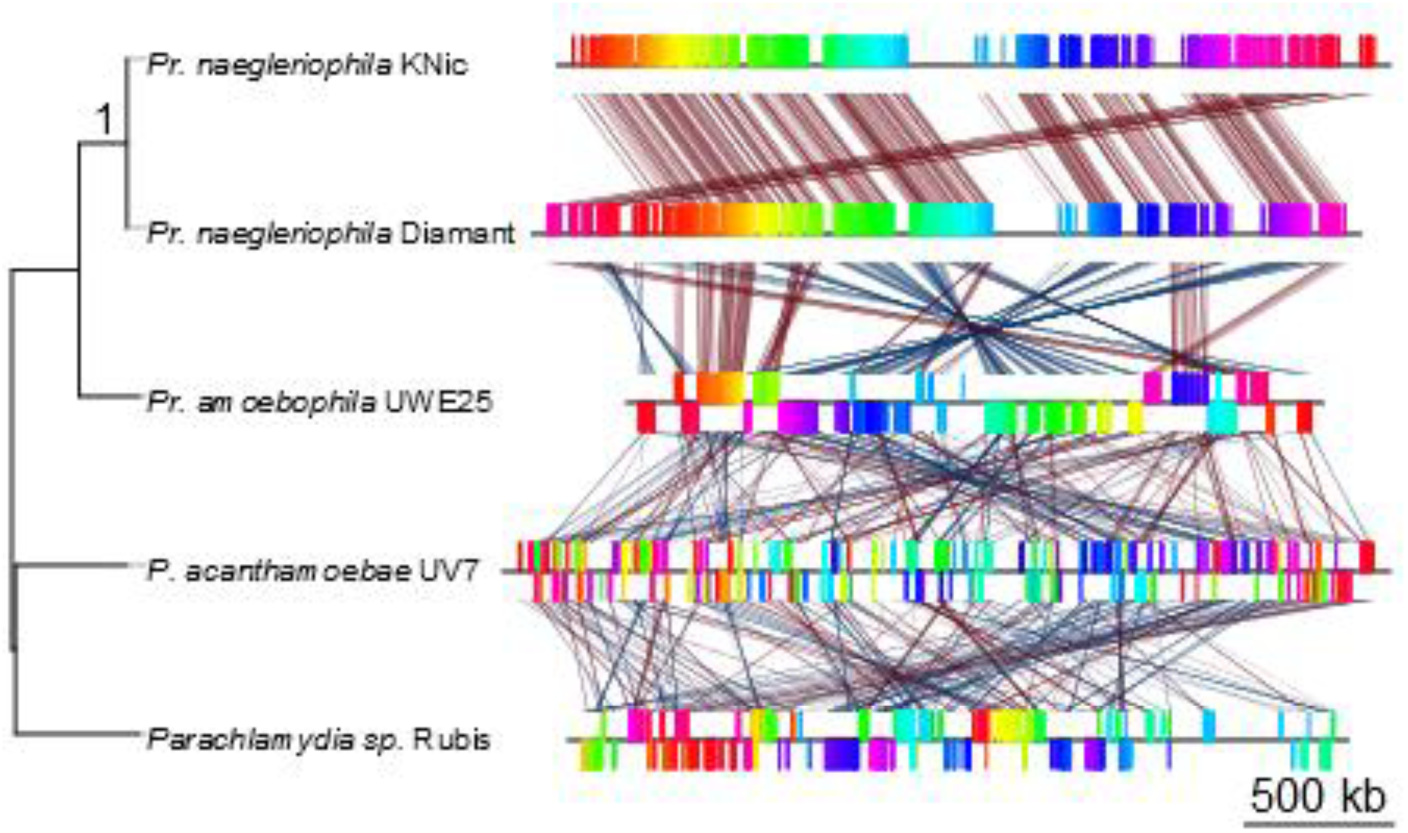
Genomic rearrangements in the *Parachlamydiaceae* family. Left side, the phylogenetic branching of bacterial strains as inferred by a Neighbor-Joining tree reconstruction based on 5 conserved proteins (DnaA, FtsK, HemL, FabI and SucA). Right side, visualization of genomic rearrangements in the family *Parachlamydiaceae.* The two strains of the species *Protochlamydia naegleriophila* are highly collinear, with no apparent rearrangement other than due to the choice of the start of the genome sequence. With increasing distances between organisms, the genomes show increasing number of rearrangements.

### pPNK is an F-type conjugative megaplasmid

The bacterial chromosome was circularized, leaving behind several contigs with a 23-fold coverage, slightly higher than the 16-fold chromosomal coverage. These contigs formed a 145,285 bp large plasmid - the largest known plasmid in the order *Chlamydiales.* The plasmid pPNK presents a GC content of 37.2% and includes 160 genes among which are several transposase and integrase remnants, doc proteins, and systems for the maintenance of the plasmid (*parA* and PNK_p0119) that are all characteristic of extra-chromosomal elements.

The plasmid also encodes a type IV secretion system with highest similarity to the F-type system found in the genomic island of *Pr. amoebophila* UWE25 (Greub et al. 2004), in the plasmid of *Chlamydia* sp. Rubis (hereafter named *Parachlamydia sp.* Rubis) and *S. negevensis* (Collingro et al. 2011) and to the remnants *traU, traN* and *traF* present in members of the family *Parachlamydiaceae* (Greub et al. 2009; Collingro et al. 2011) (**Figure 2**). The type IV secretion system of *Parachlamydia* sp. Rubis is located on a 30 kb long contig that should be circularized as a small plasmid to retain the colinearity of the *tra* operon with other bacteria. This small plasmid would therefore contain almost exclusively the *tra* operon as well as core genes for plasmid replication such as *parA. Parachlamydia* sp. Rubis and KNic *tra* operons share a striking colinearity. The comparison of gene conservation shows that *traN* has undergone different rearrangements in both *Pr. amoebophila* strains, and *traC* was split in strain R18. On the other hand, *Parachlamydia sp.* Rubis, *S.* negevensis and *Pr. naegleriophila* KNic, the three bacteria that possess the *tra* operon on a plasmid, retained intact genes. Moreover, these bacteria present Ti-type *traA* and *traD* genes downstream that share similarity to and other amoeba-infecting bacteria such as *Rickettsia bellii* and *Legionella* spp.

**Figure 2.**
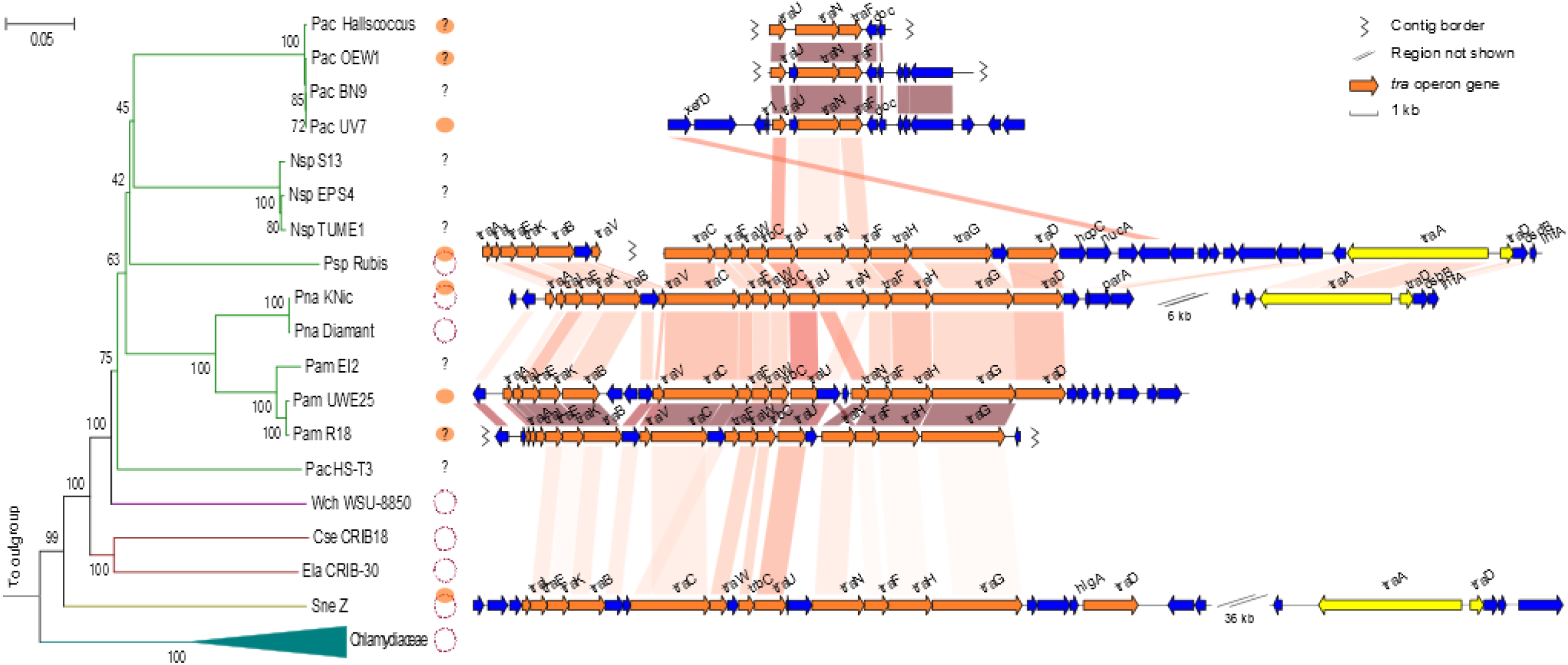
*Chlamydiales* order, plasmids and type IV secretion system. The left panel represents a neighbor joining tree of bacteria belonging the order *Chlamydiales* whose genome sequences is available based on four conserved proteins (DnaA, FtsK, HemL, FabI). The presence of a plasmid in each strain is represented by a small circular DNA molecule, and the draft genomes with no known plasmid described are indicated by a question mark as plasmids may be hidden among the numerous contigs. Orange ovals indicate the presence of a conjugative *tra* operon on the plasmid or in the bacterial chromosome. The right panel shows the conservation of the type IV secretion system *tra* operon and the surrounding genes. Pac: *P. acanthamoebae,* Nsp: *Neochlamydia* sp., Psp: *Protochlamydia sp.,* Pna: *Pr. naegleriophila,* Pam: *Pr. amoebophila,* PacHS-T3: *Parachlamydiaceae* bacterium HS-T3.

A CRISPR –Cas system for the first time within *Chlamydiales*

In *Pr. naegleriophila,* the CRISPR locus comprises eight 28bp-long repeats separated by 33bp-long spacers. The upstream operon of CRISPR-associated genes from the *E. coli* subtype I-E consists of the core genes *cas1-2,* the type I gene *cas3* and subtype-specific genes *cse1-2, cas5, cas6e* and *cas7* (**Figure 3**). An almost identical cas operon and a CRISPR locus were identified in *Pr. naegleriophila* ‘Diamant’ (**Figure 3**). On the contrary, this system is absent from other *Parachlamydiaceae* such as strains *Pr. amoebophila* UWE25, EI2 and R18. Although a confirmed CRISPR locus is predicted by CRISPRfinder (Grissa et al. 2007) in the recently released genomes of *Neochlamydia sp.* (Domman et al. 2014; Ishida et al. 2014), no *cas* genes could be identified and the repeats were found to be due to a highly repeated protein sequence.

**Figure 3.**
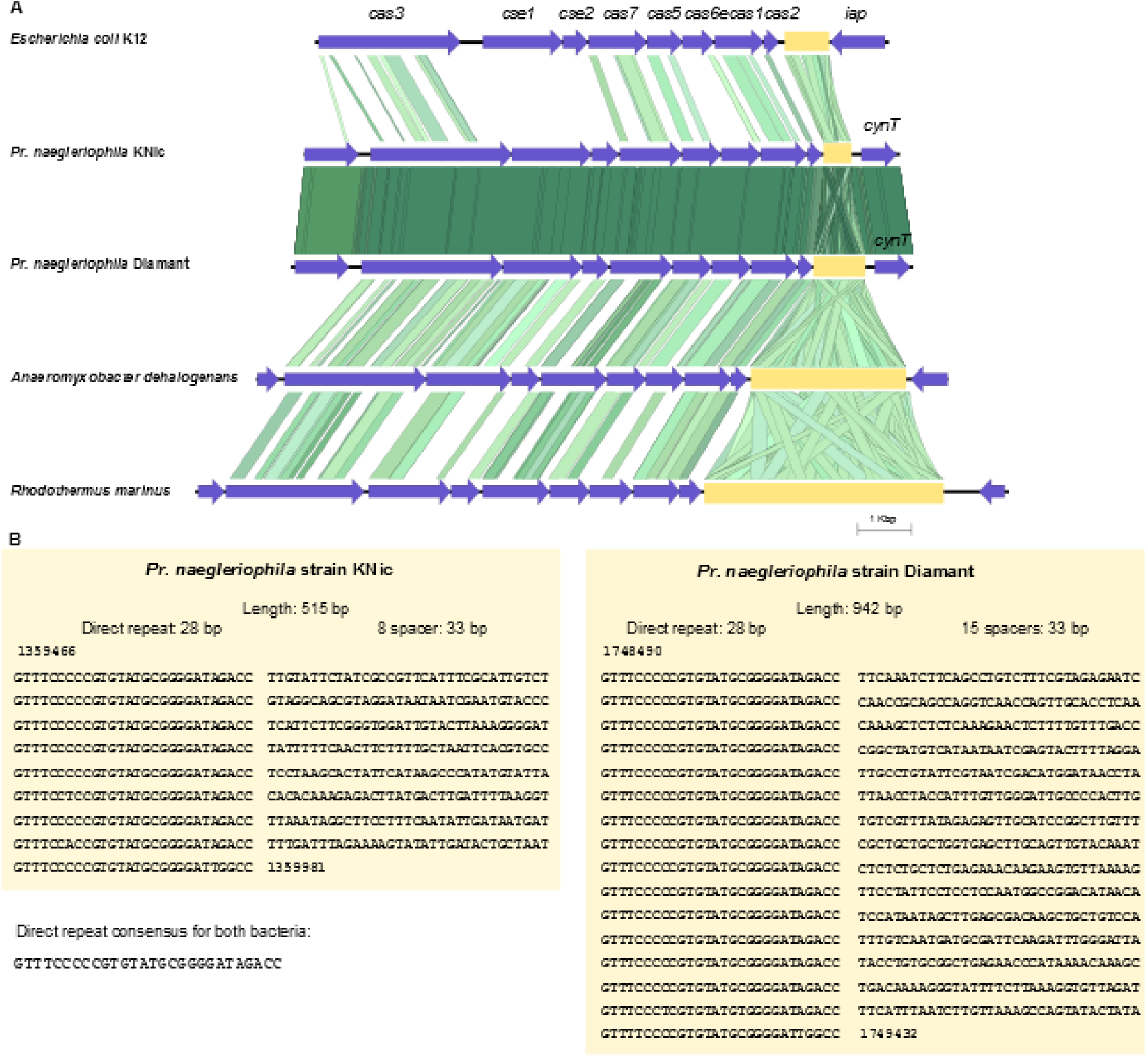
CRISPR locus and its associated genes. A. CRISPR associated genes (CAS) consist of eight coding sequences, *cas3, cse1, cse2, cas7, cas5, cas6e, cas1* and *cas2,* shown in blue within their genomic environment. Green lines connecting the genes in different organisms represent BLAST sequence homology with a gradient from light green to dark green for low to high percentage sequence identity, respectively. Genes neighboring the CRISPR locus present homology in *Pr. naegleriophila* genomes, but not to other genomes showing that the site of CRISPR locus insertion in *Pr. naegleriophila* genomes is different than in other bacteria. CRISPR repeats are found directly downstream of the CAS operon, as highlighted by the yellow box. B. Direct repeats and spacer sequences are detailed below.

The CRISPR spacers could give an interesting imprint of recent invasions by extrachromosomal elements, but unfortunately no significant homology was found by BLASTN against the non-redundant nucleotide database (nt) for strains KNic and Diamant (Sup. table 1 and sup. table 2). Genes surrounding this locus are found in conserved order in all *Protochlamydia* species indicating that this CRISPR region has most likely been acquired by horizontal gene transfer after the divergence of *Pr. naegleriophila* from *Pr. amoebophila*. The gene operon structure is commonly found in bacteria and two species present particular homology to *Pr. naegleriophila* KNic CRISPR locus: *Anaeromyxobacter dehalogenans,* a *Deltaproteobacteria* from soil and *Rhodothermus marinus,* a *Bacteroidetes* (**Figure 3**).

## DISCUSSION

The genome sequence of *Pr. naegleriophila* presented in this contribution permitted to grasp initial hints on mechanisms triggering the evolution of *Protochlamydia.* We could evidence the presence of a CRISPR-locus in the chromosome for the first time in the order *Chlamydiales.* The sequencing data also revealed the presence of a plasmid that encodes a type IV secretion system and is only partially similar to the genomic island of *Pr. amoebophila.*

Amoebae were proposed to act as a reservoir of different amoebae-resisting bacteria where horizontal gene transfer may preferentially take place (Moliner et al. 2010). The presence of a an F-like conjugation plasmid putatively involved in DNA transfer in *Pr. naegleriophila* stresses the likelihood of gene exchange with other bacteria or with the eukaryotic host. The maintenance of intact *tra* genes in bacteria possessing the *tra* operon on a plasmid, suggest that the system has retained functionality, whereas it has evolved towards pseudogenisation and deletion after being integrated in the genome of *Pr. amoebophila* strains and *P. acanthamoebae* strains.

The presence of an F-type conjugative operon in the plasmid or in the chromosome of various strains combined with the lack of conjugative operon in the plasmid or in the chromosome of the *Waddliaceae, Criblamydiaceae* (Bertelli et al. 2015, 2014) and some *Parachlamydiaceae* challenges the most parsimonious scenario proposed by Collingro *et al* (Collingro et al. 2011) that plasmids evolved from a single conjugative plasmid acquired by an ancestor of the *Parachlamydiaceae, Waddliaceae,* and *Simkaniaceae.* In favor of this hypothesis is the shared presence of the Ti-type *traA* and *traD* in the paraphyletic *Parachlamydia sp.* Rubis, *Pr. naegleriophila* KNic, as well as *S. negevensis.* If this hypothesis is correct, the plasmid and its *tra* operon were integrated within the chromosome at least twice in the genus *Parachlamydia* and *Protochlamydia.* The *tra* operon was completely lost several times, in the families *Waddliaceae* and *Criblamydiaceae,* in the genus *Neochlamydia,* and in some strains of *Protochlamydia* and *Parachlamydia* (**Figure 2**). Furthermore, it has been partially lost in the *Parachlamydia* genus, where only a few genes remain. An alternative scenario of separate acquisition of *tra* operon by an ancestor of *Simkania* and members of the family *Parachlamydiaceae* may be envisioned. In any case, this highlights the highly dynamic nature of the *Chlamydia-related* genomes and the potential of the *tra* operon to be readily transferred, acquired, and lost among these bacteria.

A CRISPR-Cas system has been reported in approximately 50% of bacteria with sporadic distribution patterns suggesting that CRISPR loci are subject to frequent horizontal gene transfer, a hypothesis supported by the presence of CRISPR loci on plasmids (Haft et al. 2005). The CRISPR locus of *Pr. naegleriophila* and its associated genes have most probably been acquired horizontally but the proteins have insufficient homology to infer a direct transfer from a given organism. This CRISPR-Cas system is of a different subtype than that of another intracellular amoeba-resisting bacteria *F. novicida* ruling out the possibility of intra-amoebal transfer between these organisms. The functionality and the exact role of this CRISPR-Cas system in *Pr. naegleriophila* remains to be determined by laboratory experiments but by similarity to the type IE locus present in *E. coli,* we can hypothesize that it plays a role in preventing DNA acquisition or protecting against phages. Although *Pr. naegleriophila* is an obligate intracellular bacteria, it may still be exposed to phages similarly to other *Chlamydia* species (Śliwa-Dominiak et al. 2013). The difference in CRISPR spacers between *Pr. naegleriophila* strains KNic and Diamant clearly highlights the dynamic and likely functional status of the system, as well as the exposure of such obligate intracellular bacteria to DNA of foreign origin. The absence of similarity between CRISPR spacers and sequences of the non-redundant nucleotide database underlines the currently limited knowledge on phages and extrachromosomal DNA circulating in amoebae-resisting bacteria, especially those growing in the ubiquitous amoeba *Naegleria.*

Based on the complete genome sequence of *P. acanthamoebae* UV-7 and *Pr. amoebophila* UWE25 as well as four draft genomes, Dommann *et al.* (Domman et al. 2014) suggested the occurrence of few rearrangements within genera of the family *Parachlamydiaceae*. Beyond the difficulty to make such an assessment based on the alignment of highly fragmented draft genome sequences available in public databases, the genomes used in the latter study are different strains of the same species and might therefore not reflect genome evolution at the genus level. Indeed, our comparison shows the absence of rearrangements between the two *Pr. naegleriophila* strains KNic and Diamant, but an increasing number of genome rearrangements with further distantly-related organisms. This highlights the need for complete genomes to precisely unravel the bacterial evolution occurring by recombination. The complete genome sequence of *Pr. naegleriophila* represents a first step toward the understanding of mechanisms triggering genome evolution and evolutionary pressures at play in the *Parachlamydiaceae* family.

## MATERIALS AND METHODS

### Culture and purification of *Pr. naegleriophila*

*Protochlamydia naegleriophila* strain KNic was grown in *Acanthamoeba castellanii* ATCC 30010 at 32°C using 75 cm^2^ cell culture flasks (Becton Dickinson, Franklin Lakes, USA) with 30 ml of peptone-yeast extract glucose broth. *Pr. naegleriophila* were purified from amoebae by a first centrifugation step at 120 × *g* for 10 min. Then, remnants from amoebae were removed from the re-suspended bacterial pellet by centrifugation at 6500 × *g* for 30 min onto 25% sucrose (Sigma Aldrich, St Louis, USA) and finally at 32000 × *g* for 70 min onto a discontinuous Gastrographin (Bayer Schering Pharma, Zurich, Switzerland) gradient (48%/36%/28%).

### Genome sequencing, assembly and gap closure

*Pr. naegleriophila* genomic DNA was isolated with the Wizard Genomic DNA purification kit (Promega Corporation, Madison, USA). Reads obtained with Genome Sequencer 20^TM^ (Droege and Hill 2008) by Roche Applied Science (Penzberg, Germany) were assembled using Newbler V1.1.02.15 yielding 93 large contigs with a mean 16x coverage. Scaffolding on *Pr. amoebophila* strain UWE25 and PCR-based techniques were used to close the gaps between those contigs. Solexa 35 bp reads obtained from sequencing with Genome Analyzer GaIIx (Bennett 2004) by Fasteris (Plan les Ouates, Switzerland) were then mapped to the final assembly with BWA (Li & Durbin 2009) and visualized with Consed (Gordon & Green 2013). Homopolymer errors were corrected in the plasmid and the chromosome sequence after manual inspection of discrepancies covered by >2 reads with a phred quality score of the base >10. Sequence start was placed in an intergenic region closest to the minimum of the GC skew, as determined with a sliding window of 100nt.

### Genome annotation

GenDB 2.4 pipeline (Meyer et al. 2003) was used for a first automatic annotation of the genome that was followed by manual curation of annotation. Coding sequence (CDS) prediction was performed using Prodigal (Hyatt et al. 2010). All predicted CDS were submitted to similarity searches against nr, Swissprot, InterPro, Pfam, TIGRfam and KEGG databases. Putative signal peptides, transmembrane helices and nucleic acid binding domains were predicted using respectively SignalP (Petersen et al. 2011), TMHMM (Krogh et al. 2001) and Helix-Turn-Helix (Dodd & Egan 1990). Genome annotation was manually curated with a scheme as proposed in (Bertelli et al. 2015). The complete and annotated genome sequences have been deposited in the European Nucleic Archive under the project PRJEB7990 with accession numbers LN879502 and LN879503.

### Genome analysis

To identify CRISPR repeats, the genome sequences were submitted to CRISPRFinder (Grissa et al. 2007). The spacers within CRISPR locus of *Pr. naegleriophila* strains KNic and Diamant were submitted to BLASTN (Altschul et al. 1997) homology searches against the non-redundant nucleotide database. For phylogenetic reconstruction, multiple sequence alignments were performed with Muscle V3.7 (Edgar 2004), and a neighbor-joining tree was reconstructed using Mega 6 (Tamura et al. 2013) with 1000 bootstrap, poisson distribution, gamma equal to 1. The two nearly complete genomes of *Chlamydia* sp. Rubis and *Pr. naegleriophia* Diamant were reordered with Mauve (Darling et al. 2004) by similarity to the closest available complete genome sequence *P. acanthamoebae* UV7 and *Pr. naegleriophila* KNic, respectively. These genomes and the complete genomes were aligned using Mauve and the alignment was represented using GenoPlotR (Guy et al. 2010). Genomic islands were predicted using IslandViewer (Dhillon et al. 2015). Home-made scripts for data analysis and visualization were written in R (Cran 2010).

## ACKNOWLEDGMENTS

We are grateful to Sébastien Aeby (University of Lausanne, Switzerland) for his technical help during the gap closure stage. We would like to thank Burkhard Linke (Justus-Liebig-University Giessen, Germany) for his assistance in maintaining this GenDB project. We acknowledge technical assistance by the Bioinformatics Core Facility at JLU Giessen and access to resources financially supported by the BMBF grant FKZ 031A533 within the de.NBI network. Part of the computations was performed at the Vital-IT (http://www.vital-it.ch) Center for high-performance computing of the SIB Swiss Institute of Bioinformatics.

## SUPPLEMENTARY FIGURES

**Figure S1.**
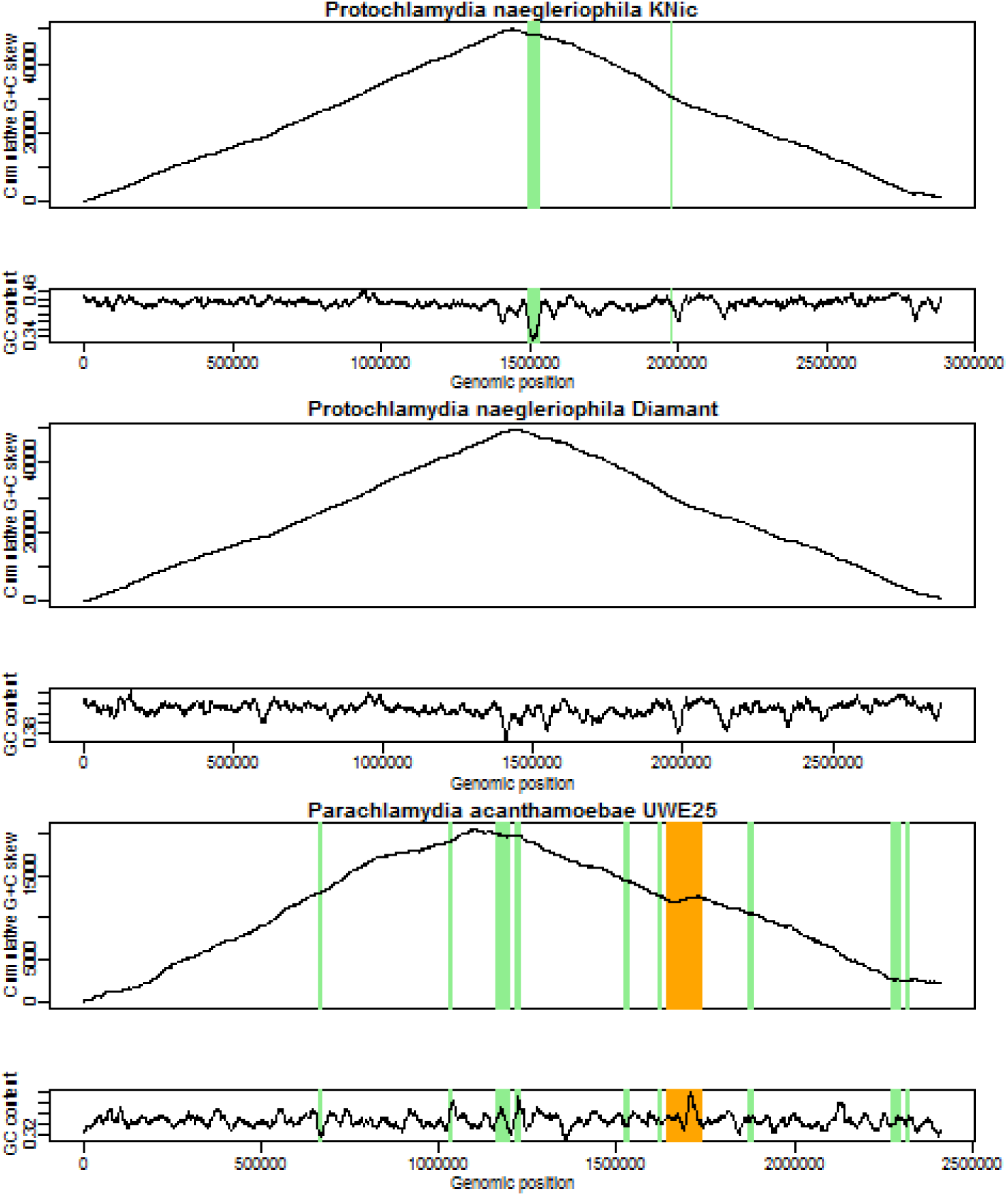
Nucleotide biases and genomic islands. The GC content and the bias of G versus C (GC skew) were determined in a sliding window of 1000 bp along the genome sequence of *Pr. naegleriophila* KNic, *Pr. naegleriophila* Diamant and *Pr. amoebophila* UWE25. The maximum of the GC skew indicates the putative terminus of replication. The origin of replication is found at the minimum of the curve and has been used to assess the first base of the genomic sequences. The contigs of *Pr. naegleriophila* Diamant have been reordered by similarity to strain KNic and the putative plasmid removed. The location of predicted GIs for *Pr. naegleriophila* KNic and *Pr amoebophila* UWE25 is indicated in green, whereas the confirmed *Pr. amoebophila* genomic island Pam100G is indicated in orange.

**Figure S2.**
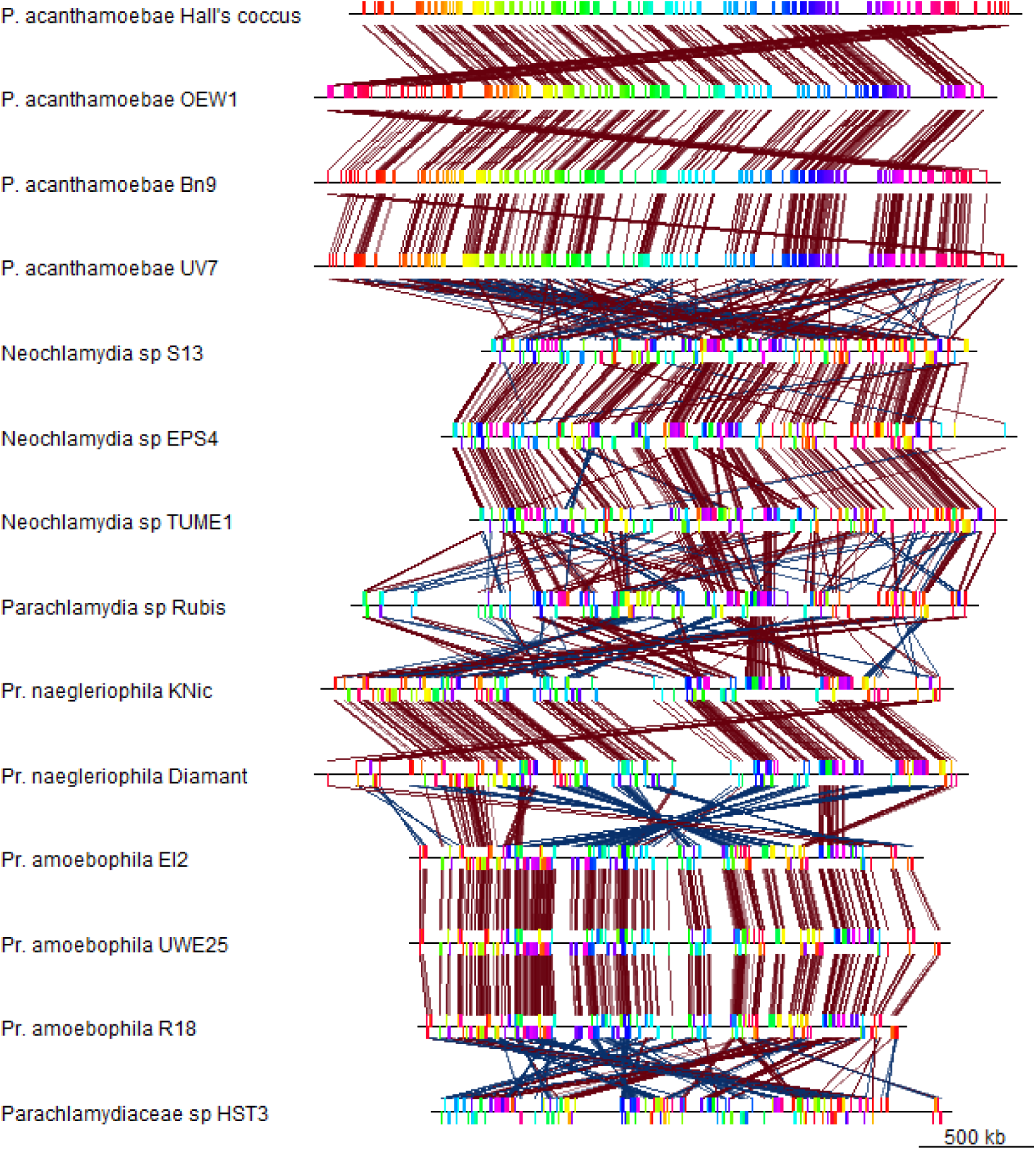
Genome rearrangements in the *Parachlamydiaceae*. This figure shows the difficulty of using draft genomes to observe genomic rearrangements. Available draft genomes of members of the family *Parachlamydiaceae* were reordered using Mauve against the complete genome of the most closely-related organism: *Pr. naegleriophila* Diamant and *Chlamydia* sp. Rubis against *Pr. naegleriophila* KNic; *Pr. amoebophila* EI2 and R18 against *Pr. amoebophila* UWE25; *P. acanthamoebae* BN9, Hall’s coccus, and OEW1 against *P. acanthamoebae* UV7; and *Parachlamydia* sp. HST3 and *Neochlamydia* sp. S13 against *Chlamydia* sp. Rubis, and *Neochlamydia* sp. EPS4 and TUME1 against *Neochlamydia* sp. S13. As expected, no rearrangement can be observed after the rearrangement of a fragmented draft genome against closely-related strain of the same species. In the case of *Neochlamydia* sp. S13 and *Parachlamydia* sp. HST3 for which no complete genome is available, numerous rearrangements can be seen, most likely as an artifact due to the reduced similarity at nucleotide level between the reordered genome and the reference genome.

## SUPPLEMENTARY TABLE

**Table S1.**
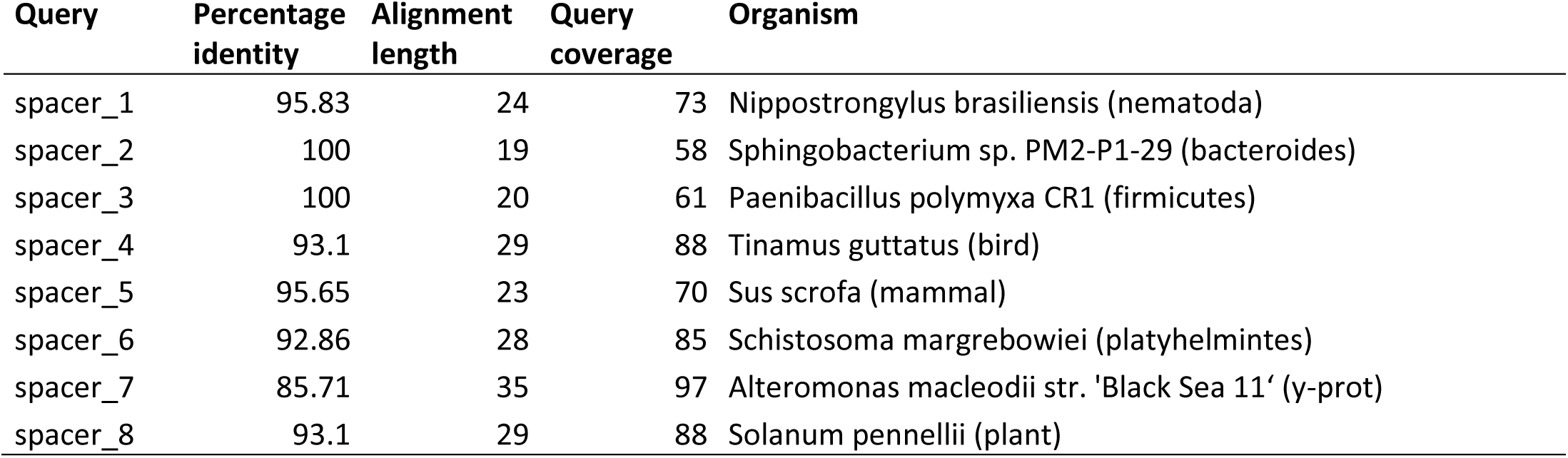
Homology of *P. naegleriophila* KNic spacers against the non-redundant nucleotide database by BLASTN

**Table S2.** Homology of *P. naegleriophila* Diamant spacers against the non-redundant nucleotide database by BLASTN

| Query | Percentage identity | Alignment length | Query coverage | Organism |
| --- | --- | --- | --- | --- |
| spacer_1 | 100 | 19 | 58 | uncultured archaeon |
| spacer_2 | 100 | 21 | 64 | Pseudomonas aeruginosa |
| spacer_3 | 90 | 30 | 91 | Populus euphratica |
| spacer_4 | 89.29 | 28 | 85 | Spirometra erinaceieuropaei |
| spacer_5 | 92 | 25 | 76 | Toxocara canis |
| spacer_6 | 95.65 | 23 | 70 | Xenorhabdus nematophila AN6/1 |
| spacer_7 | 100 | 19 | 58 | Heterobasidion irregulare TC 32-1 |
| spacer_8 | 87.5 | 32 | 97 | Kluyveromyces marxianus |
| spacer_9 | 96 | 25 | 76 | Homo sapiens |
| spacer_10 | 95.83 | 24 | 73 | Phoenix dactylifera |
| spacer_10 | 100 | 21 | 64 | Candidatus Nitrososphaera evergladensis SR1 |
| spacer_11 | 95.83 | 24 | 73 | Solanum tuberosum |
| spacer_12 | 96.3 | 27 | 82 | Staphylococcus aureus subsp. aureus MSHR1132 |
| spacer_13 | 100 | 21 | 64 | Vibrio fischeri MJ11 |
| spacer_14 | 92.31 | 26 | 79 | Borrelia garinii PBi |
| spacer_15 | 92.59 | 27 | 82 | Echinostoma caproni |

